# Distinctive nuclear zone for RAD51-mediated homologous recombinational DNA repair

**DOI:** 10.1101/2021.11.29.470307

**Authors:** Yasunori Horikoshi, Hiroki Shima, Jiying Sun, Wataru Kobayashi, Volker J. Schmid, Hiroshi Ochiai, Lin Shi, Atsuhiko Fukuto, Yasuha Kinugasa, Hitoshi Kurumizaka, Tsuyoshi Ikura, Yolanda Markaki, Shin-ichi Tate, Kazuhiko Igarashi, Thomas Cremer, Satoshi Tashiro

**Affiliations:** Department of Cellular Biology, Research Institute for Radiation Biology and Medicine, Hiroshima University, Hiroshima, Japan; Research Center for Mathematics of Chromatin Live Dynamics, Hiroshima University, Higashi-Hiroshima, Japan; Core Research for Organelle Diseases in Hiroshima University, Hiroshima, Japan; Department of Biochemistry, Tohoku University Graduate School of Medicine, Sendai, Japan; Laboratory of Chromatin Structure and Function, Institute for Quantitative Biosciences, The University of Tokyo, Tokyo, Japan; Laboratory of Structural Biology, Graduate School of Advanced Science & Engineering, Waseda University, Tokyo, Japan; Bayesian Imaging and Spatial Statistics Group, Department of Statistics, Ludwig-Maximilians-Universität München, München, Germany; Department of Mathematical and Life Sciences, Graduate School of the Integrated Sciences for Life, Hiroshima University, Higashi-Hiroshima, Japan; Institute of Medical Imaging and Digital Medicine, School of Medical Imaging, Xuzhou Medical University, Xuzhou, China; Department of Radiology, Affiliated Hospital of Xuzhou Medical University, Xuzhou, China; Department of Ophthalmology and Visual Sciences, Graduate School of Biomedical Sciences, Hiroshima University, Hiroshima University, Hiroshima, Japan; Laboratory of Chromatin Regulatory Network, Department of Genome Biology, Radiation Biology Center, Graduate School of Biostudies, Kyoto University, Kyoto, Japan; Division of Anthropology and Human Genomics, Biocenter, Ludwig-Maximilians University Munich, Grosshadernerstrasse 2, 82152 Planegg-Martinsried, Germany; Department of Biological Chemistry at the David Geffen School of Medicine, University of California Los Angeles, Los Angeles, CA, USA; Center for Regulatory Epigenome and Diseases, Tohoku University Graduate School of Medicine, Sendai, Japan

**Author notes:** MPI of Biochemistry, Am Klopferspitz 18, 82152 Martinsried, Germany.

## Abstract

Genome-based functions are inseparable from the dynamic higher-order architecture of the cell nucleus. In this context, the repair of DNA damage is coordinated by precise spatiotemporal controls that target and regulate the repair machinery required to maintain genome integrity. However, the mechanisms that pair damaged DNA with intact template for repair by homologous recombination (HR) without illegitimate recombination remain unclear. This report highlights the intimate relationship between nuclear architecture and HR in mammalian cells. RAD51, the key recombinase of HR, forms spherical foci in S/G_2_ phases spontaneously. Using super-resolution microscopy, we show that following induction of DNA double-strand breaks RAD51 foci at damaged sites elongate to bridge between intact and damaged sister chromatids; this assembly occurs within bundle-shaped distinctive nuclear zones, requires interactions of RAD51 with various factors, and precedes ATP-dependent events involved the recombination of intact and damaged DNA. We observed a time-dependent transfer of single-stranded DNA overhangs, generated during HR, into such zones. Our observations suggest that RAD51-mediated homologous pairing during HR takes place within the distinctive nuclear zones to execute appropriate recombination.

## Introduction

During the last two decades compelling evidence has accumulated that transcription and DNA/chromatin replication are inseparably intertwined with an evolutionary conserved, exceedingly complex and dynamic structural nuclear landscape. Microscopic and biochemical evidence has demonstrated that in mammalian nuclei this landscape is built up from chromosome territories (CTs) with chromatin domains (CDs)/topologically associated domains (TADs) (Cremer, Cremer et al., 2020, Dekker, Belmont et al., 2017). In contrast, little is known about the roles played by chromatin and nuclear architecture in DNA repair (Lukas, Lukas et al., 2011, Misteli & Soutoglou, 2009).

Homologous recombinational repair (HR) is a versatile repair system for DNA damage, such as DNA double-strand breaks (DSBs) and stalled replication forks. Briefly, after the generation of a DSB, histone H2AX phosphorylated at serine 139, called γH2AX (Rogakou, Boon et al., 1999), forms nuclear foci near the site of damage, followed by an accumulation of the Replication protein A (RPA) complex. Under these conditions subunits of the RPA complex are phosphorylated (Binz, Sheehan et al., 2004); for example, the 32 kDa subunit (RPA2) becomes phosphorylated at serine 33. These molecules coat the resected single-stranded DNAs (ssDNAs) formed at the DSBs; the homology search occurs once RPA is replaced by RAD51, a key player in HR (Renkawitz, Lademann et al., 2014).

HR requires the pairing of a damaged DNA sequence with its intact, homologous counterpart. The actual motions and dynamic structural changes of chromatin required to bring damaged and intact sequences present in sister CDs in direct contact prior to pairing are currently unknown, in stark contrast to the wealth of information with respect to the molecular features of HR. In case of a DSB formed in a repetitive sequence, an intact homologous sequence may be located nearby in the same damaged CD. However, HR of a DSB generated within a single copy sequence requires a much more complex structural reorganization. In mammalian cell nuclei (diameter ca. 10 μm) homologous CTs are often widely separated (Bolzer, Kreth et al., 2005, Stevens, Lando et al., 2017). Such large distance movements are essentially involved in homologous chromosome pairing during meiotic prophase (Scherthan, 1996). In somatic cell nuclei the elaborate mechanism for homologous pairing, which has evolved for meiosis, does not exist, and to which extent CTs can undergo large scale movements in most somatic cell types, is still a controversial issue. Although the major repositioning of CTs during DNA damage-repair response was suggested, no evidence for increased homologous associations was reported (Mehta & Haber, 2014). Structural constraints provided by higher-order nuclear architecture may explain why HR in mammalian cell nuclei is typically restricted to the S- and G2-phases of the cell cycle, when sister CDs become available, which are typically located up to several hundred nanometers apart from each other (Stanyte, Nuebler et al., 2018). This implies that entire sister CDs or chromatin loops must overcome such distances for homolog pairing.

In the present study, we employed 3D imaging with confocal laser scanning microscopy (CLSM) and high-resolution 3D structured illumination microscopy (3D-SIM) (Schermelleh, Ferrand et al., 2019) to study the topography of RAD51. Our findings provide evidence that nuclei assemble distinctive nuclear zones for DNA damage repair based on HR. We demonstrate that RAD51 foci elongate at sites of DSBs along such bundle-shaped zones. Our functional analysis suggests that this process requires various protein-protein interaction, but not ATPase activity of RAD51. We argue that intranuclear assemblies containing RAD51 within the zones may bridge sister chromatids and serve as a scaffold for homologous pairing of intact and damaged DNA sequences.

## Results

### Elongation of RAD51 foci upon DNA damages

A modest number of RAD51 foci can be observed in unperturbed nuclei (Tashiro, Kotomura et al., 1996). However, upon challenging cells with DNA-damaging agents, the number of foci increases dramatically (Haaf, Golub et al., 1995, Tashiro, Walter et al., 2000). To examine the difference between these foci, we analyzed 3D images of X-ray-irradiated or unirradiated cell nuclei using 3D-SIM (Fig 1A). After two hours of ionizing radiation, we observed a significant change in the shape of these foci: they became elongated and ellipsoidal (Fig 1B and C). Furthermore, the RAD51 foci closely associated with γH2AX foci, were significantly more elongated than those that were not near γH2AX (Fig 1C).

**Figure 1.**
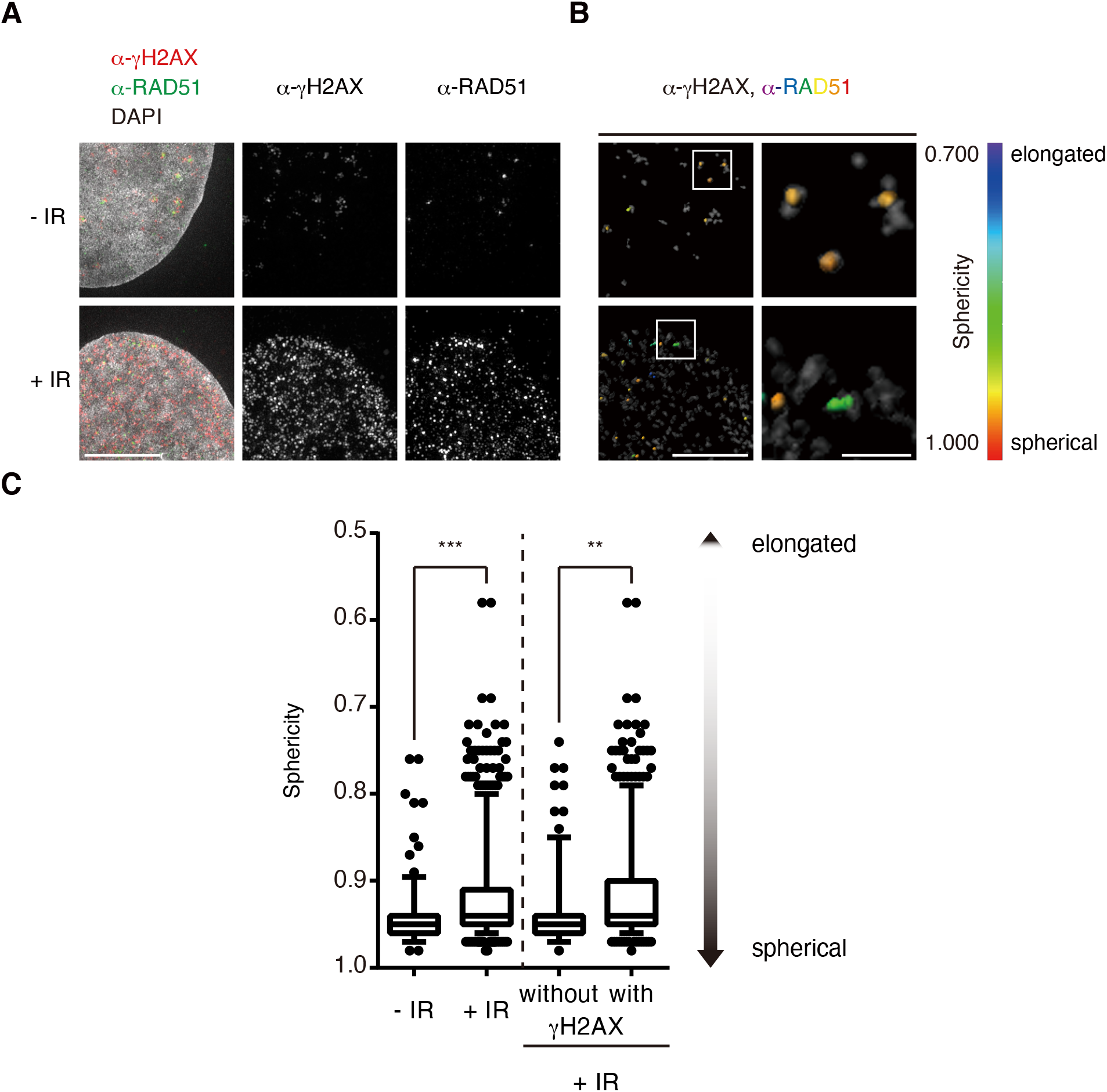
DNA damage-dependent elongation of RAD51 foci. A Before and at 2 hours after exposure to 2 Gy X-ray, asynchronous GM0637 cells were pre-treated with CSK buffer and then fixed. Endogenous RAD51 and γH2AX were detected by immunofluorescence and DNA was stained with DAPI. Z-stack images were acquired by 3D-SIM and representative projection images are shown. In merged images, RAD51, γH2AX and DNA were visualized in green, red and grayscale, respectively. Scale bar: 5 μm. B The 3D reconstruction image of (A). Each RAD51 focus was colored according to its respective sphericity, and γH2AX was visualized in grayscale. Scale bars: 5 μm or 1 μm (insets). C Statistical analysis of the sphericities of the respective RAD51 foci in cell nuclei under indicated experimental conditions. Data is presented as a Tukey box-and-whiskers plot, with values that are above and below the whiskers drawn as individual dots. ***P<0.0001, Mann–Whitney-Wilcoxon test.

We considered two possibilities for the role of the spherical foci in undamaged nuclei: some may serve as storage centers for repair factors, while others may be involved in the repair of intrinsic or spontaneous DNA breaks (Tubbs & Nussenzweig, 2017), which often occurs at stalled replication forks (Branzei & Foiani, 2010). To test this, we analyzed nuclear foci in cells treated with hydroxyurea (HU), which induces replication-associated DNA damage and fork collapse (Koç, Wheeler et al., 2004). We found that the sphericity of the RAD51 foci in HU-treated cells was intermediate between that of the two other conditions (Fig EV1). Moreover, they retained their shape regardless of the duration of drug exposure. In brief, RAD51 foci associated with stalled replication forks were distinctly more spherical than those involved in IR-induced DSB repair. Taken together, our findings suggested that RAD51 foci alter their shape depending on the type of repair that is needed.

### Spatial association of RAD51 foci with damages and intact sister chromatids

Next, we employed the TALEN system (Bogdanove & Voytas, 2011) to study the topographical relationship of RAD51 foci with a DSB introduced at a specific genomic site. We targeted a DSB to the Breakpoint Cluster Region (BCR) in the MLL gene locus on 11q23 (Fig EV2), one of the most common translocation breakpoint sites in mixed-lineage leukemias (Krivtsov & Armstrong, 2007). We then performed two-color fluorescence in situ hybridization on structurally preserved nuclei (3D-FISH) using DNA probes that flank the DSB (green and red, respectively), in combination with immunofluorescence staining of RAD51. In principle, sister chromatid HR takes place after DNA replication. Here, we took advantage of DNA FISH analysis to eliminate G_1_ phase cell nuclei, without doublets of red and/or green signals (Fig EV3A). CLSM light optical serial sections were recorded and revealed a close spatial association of the FISH signals with RAD51 only in TALEN-treated cell nuclei (Fig EV3B and C). This is consistent with the notion that TALEN induced DSBs at BCR in the MLL locus. The efficiency of cleavage monitored by PCR was roughly 40%, and in TALEN treated cells around 40% of sister chromatids were cleaved, producing separated red and/or green spots by DNA FISH analysis (Fig EV2B, C, EV3D and E). Whereas in some nuclei green and red FISH foci were found more than 1 μm apart, in others we observed all three signals, paired red or green FISH dots and RAD51 staining together within a radius of 400 nm (Fig EV3B, D and E). In such cases the RAD51 focus was localized between a pair of like-colored FISH signals, suggesting that these RAD51 foci bridge sister chromatids harboring DSBs.

### Topography of RAD5f foci within the nuclear landscape

We next investigated the localization of RAD51 foci with respect to the higher-order folding of chromatin (Schmid, Cremer et al., 2017). In this analysis, we assigned pixels representing DAPI-stained DNA in each SIM section to seven, color-coded DAPI intensity classes (class 1 – 7). Briefly, class 1 represented the nearly DNA-free interior of the interchromatin compartment (IC) and wider lacunae; Classes 2-4 represented a zone of less compacted chromatin (presumably ‘open’ chromatin) at the periphery of chromatin domain clusters (CDCs), Class 5 was considered to be a transition zone between ‘open’ and ‘closed’ chromatin, while Classes 6 and 7 reflected the most compact chromatin within CDCs. Indeed, we confirmed the localization of SC35 or PML bodies within IC by use of this assay (Fig EV4). Although pixels assigned to RAD51 and γH2AX were present in all 7 classes, they were not randomly distributed and were significantly enriched in Class 3 to 5 (Fig 2). Of note, this distribution pattern was not altered by exposure to X-rays, although we did observe a conspicuous increase in the number of RAD51 foci after IR (Fig 2B).

**Figure 2.**
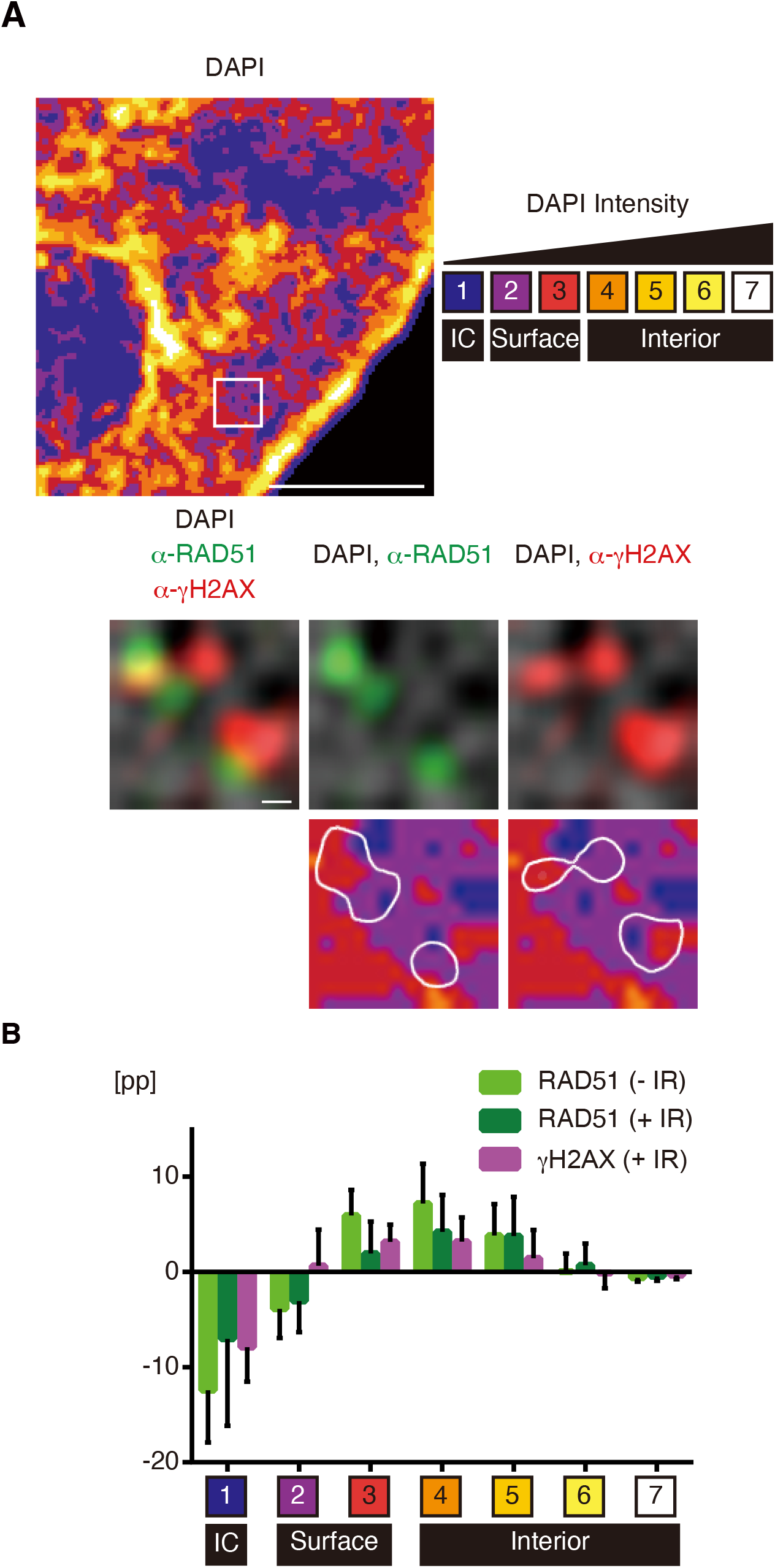
Topography of endogenous RAD51 foci within the nuclear landscape. A Before and at 2 hours after exposure to 2 Gy X-ray, asynchronous GM0637 cells were pre-treated with CSK buffer and then fixed. Endogenous RAD51 and γH2AX were detected by immunofluorescence and visualized in green and red, respectively. Z-stack images were acquired by 3D-SIM. DNA was stained with DAPI and displayed in a false-color code according to the respective signal intensities, ranging from class 1 (blue, close to background intensity) to class 7 (white, highest), or grayscale. A representative single section image is shown. Scale bars: 5 μm and 200 nm (insets). B Over/underrepresentation of RAD51 and γH2AX in DAPI intensity classes of cell nuclei (n = 10). Values represent means with SD.

### RAD51 accumulation in the distinctive nuclear zone

It has been shown that an excessive amount of RAD51 protein exhibited characteristic, elongated bundle-like structures in the nucleus independently of DNA damages (Raderschall, Bazarov et al., 2002). These findings lead us to formulate the hypothesis that RAD51 foci are formed by accumulation only within such a distinctive bundle-shaped nuclear zone.

To test this idea, we examined the time-dependent change of the shapes of RAD51 foci in living cells transiently overexpressing EGFP-fused RAD51. To confirm that the N-terminally tagged EGFP to RAD51 (EGFP-N-RAD51) fusion retains recombinase activity, we performed an *in vitro* D-loop formation assay with the overexpressed protein (Fig EV5A-D). This functionality contrasts with the C-terminal fusion of EGFP to RAD51, which is deficient for repair (Essers, Houtsmuller et al., 2002). As expected, the time-lapse imaging provided evidence that the elongated bundle-like signals resulted from fusion of elongated EGFP-N-RAD51 foci (Fig 3A). Subsequently, we generated UVA-microirradiated nuclear stripes with DSBs and then observed time-dependent changes of EGFP-N-RAD51 in the presence of DNA damages. An increase in signal intensity at the DSB-containing nuclear stripes was detected 2h after microirradiation (mIR), indicating an accumulation of RAD51 near DSBs (Fig 3B). Furthermore, we monitored a significant enrichment of a transiently overexpressed EGFP-N-RAD51 colocalizing in chromatin Classes 3 to 5 (Fig 3C and D). Likewise, these bundles appeared to span interchromatin channels to bridge CDs, consistent with the case of endogenous RAD51 (Fig 2). We also observed the responses to induction of DSBs *in vivo*, accumulation and turnover of EGFP-N-RAD51 at DNA damage sites (Fig EV5E), confirming a similar behavior of endogenous RAD51 and RAD51 with EGFP fused to its N-terminal site. More importantly, the intensity increased only in the bundle-like regions, not outside of them. That is, there was no de novo RAD51 focus formation within the microirradiated area.

**Figure 3.**
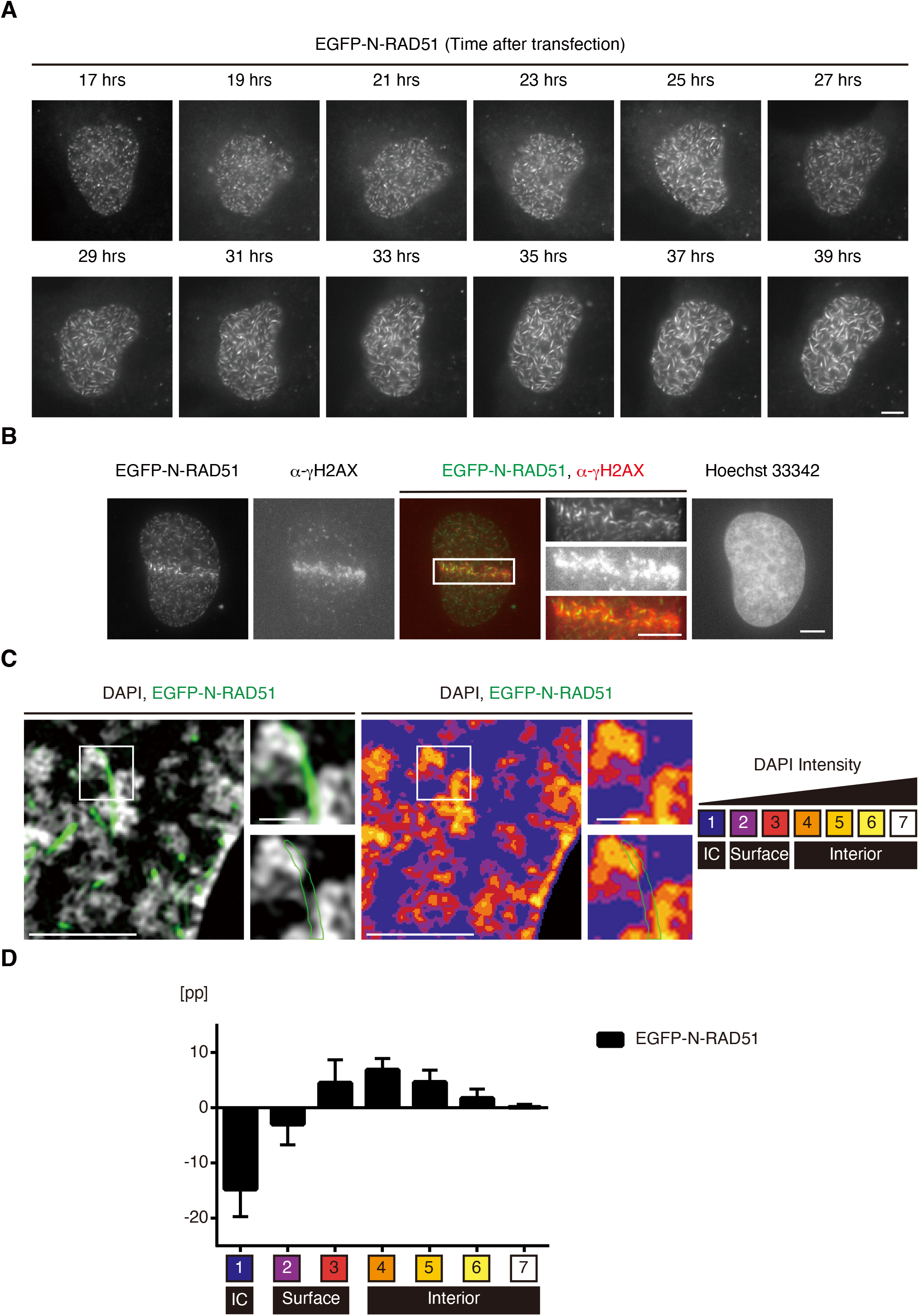
Topography of EGFP-N-RAD51 within the nuclear landscape. A Asynchronous GM0637 cells were transiently transfected with EGFP-N-RAD51 expression vector. At the indicated time points after transfection, the respective images of an identical cell nucleus were acquired by confocal laser-scanning microscopy. Scale bar: 5 μm. B Asynchronous GM0637 cells were transiently transfected with EGFP-N-RAD51 expression vector and incubated for 24 hours. Subsequently, UVA-microirradiation (mIR) was performed and fixed at 210 minutes after mIR. γH2AX was detected by immunofluorescence and DNA was stained with Hoechst 33342. Representative image acquired by wide-field microscopy is shown. In merged image, EGFP-N-RAD51 and γH2AX were visualized in green and red, respectively. Scale bar: 5 μm. C Asynchronous GM0637 cells were transiently transfected with EGFP-N-RAD51 expression vector and fixed at 24 hours after transfection. Z-stack images were acquired by 3D-SIM. EGFP-N-RAD51 was visualized in green. DNA was stained with DAPI and displayed in a false-color code according to the respective signal intensities, ranging from class 1 (blue, close to background intensity) to class 7 (white, highest), or grayscale. A representative single section image is shown. Scale bars: 5 μm and 200 nm (insets). D Over/underrepresentation of EGFP-N-RAD51 in DAPI intensity classes of nuclei (n = 16). Values represent means with SD.

### Protein-protein interactions required for accumulation of RAD51 into nuclear zones

Next, we attempted to acquire further mechanistic insights into the role of RAD51 itself in the creation of the elongated foci. Here, we investigated the nuclear distribution of four EGFP-N-RAD51 mutants that were defective in self-association (F86E), ATPase activity (G151D), BRCA2 binding (A190/192L) or interaction with SUMO (V264K), respectively (Chen, Morrical et al., 2015, Shima, Suzuki et al., 2013, Yu, Sonoda et al., 2003). As shown in Fig 4A and B, overproduced EGFP-N-RAD51-G151D exhibited a bundle-like distribution identical to wild-type, while F86E and V264K mutants failed to do so. In the case of EGFP-N-RAD51-A190/192L, although bundle-like signals were observed, they formed half as frequently as wild-type RAD51. Note that F86E mutant showed a widespread uniform nuclear distribution pattern, albeit the highest protein level among them (Fig EV6A); the expression level seems independent of this localization. We also examined the effect of a series of mutations on mobility in the nucleus. FRAP analysis of wild-type and mutant EGFP-N-RAD51 revealed a much more rapid recovery of bleached signals of mutants except EGFP-N-RAD51-G151D, as compared to wild-type (Fig 4C and D). Moreover, the increase of signal intensity at UVA-microirradiated area like wild-type was confirmed only in the cells expressing EGFP-N-RAD51-G151D (Fig 4E and F). Notably, in the case of EGFP-N-RAD51-A190/192L this behavior was not observed, even though they exhibited the bundle-like distribution. Taken together, these functional analyses suggest that three aspects of RAD51 function, self-association, interaction with SUMO and binding to BRCA2, but not ATPase hydrolysis, are responsible for the generation of elongated RAD51 foci and its subnuclear dynamics.

**Figure 4.**
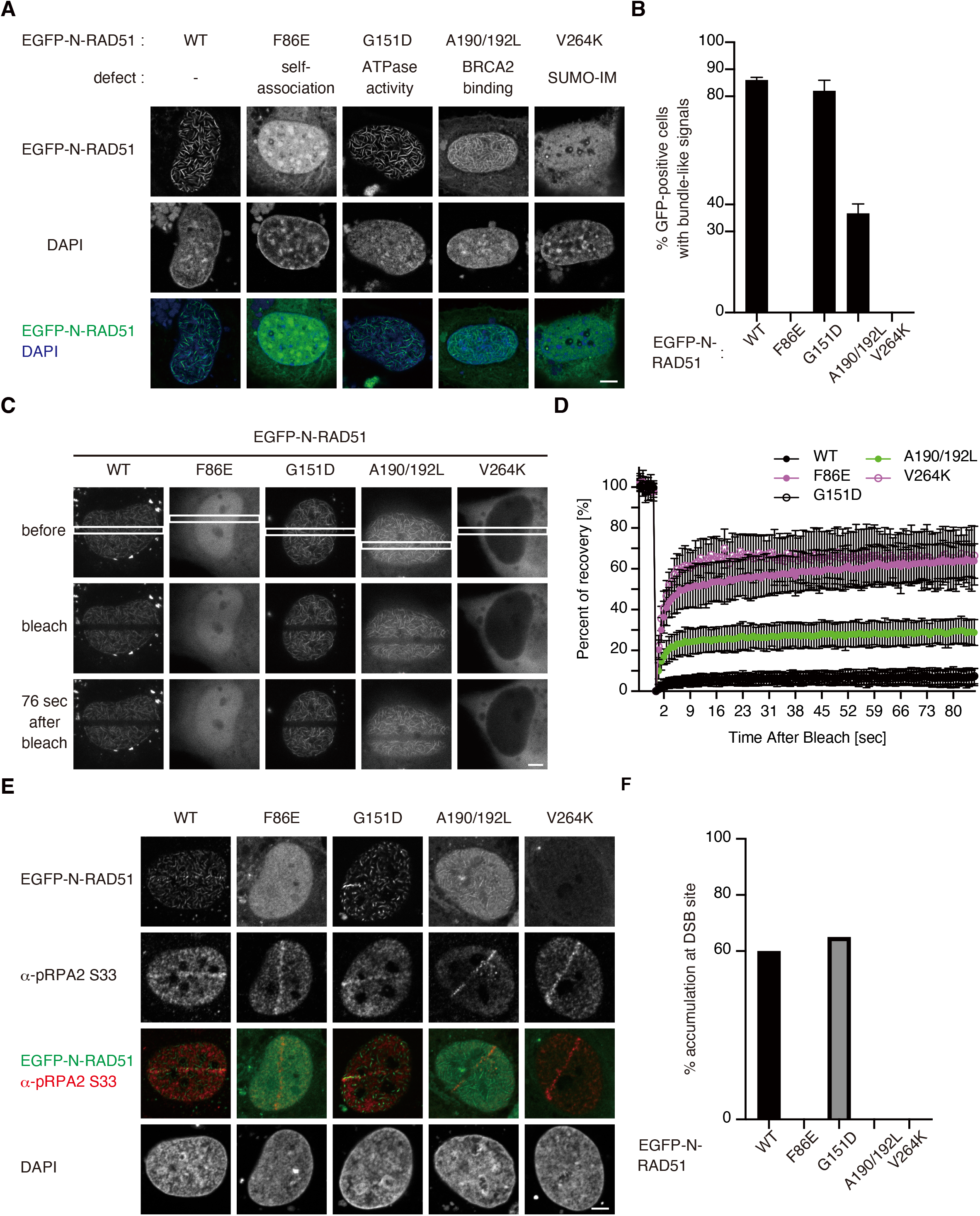
Influence of functional domains within RAD51 on its distribution and dynamics. A, B Asynchronous GM0637 cells were transiently transfected with either one of the indicated types of mutant EGFP-N-RAD51 expression vectors, defective in self-association (F86E), ATPase activity (G151D), BRCA2 binding (A190/192L) or interaction with SUMO (V264K), respectively. Cells were fixed at 24 hours after transfection and images were acquired. Representative images acquired by confocal laser-scanning microscopy are shown in (A). In merged image, EGFP-N-RAD51 and DNA were visualized in green and blue, respectively. Scale bar: 5 μm. For quantitative evaluation, every 200 cell nuclei images were also acquired by Metafer. The percentages of cells carrying the bundle-like distribution of EGFP signals are shown in (B). C, D Asynchronous GM0637 cells were transiently transfected with either one of the indicated types of mutant EGFP-N-RAD51 expression vectors. At 24 hours after transfection, fluorescence recovery after photobleaching (FRAP) analysis was performed; images were continuously acquired. Representative images are shown in (C) and enclosed areas in the top panel were bleached. Scale bar: 5 μm. The percentages of signal intensity recovery were calculated and indicated in (D). Values represent means with SDs (n = more than 20, respectively). E, F Asynchronous GM0637 cells were transiently transfected with either one of the indicated types of mutant EGFP-N-RAD51 expression vectors and incubated for 24 hours. Subsequently, UVA-microirradiation (mIR) was performed and fixed at 120 minutes after mIR. Phosphorylated RPA2 (S33) (pRPA2 S33) was detected by immunofluorescence and DNA was stained with DAPI. Representative images acquired by confocal laser-scanning microscopy are shown in (E). In merged images, EGFP-N-RAD51 and pRPA2 S33 were visualized in green and red, respectively. Scale bar: 5 μm. Each ratio of cells exhibiting the accumulation of EGFP signals at UVA-irradiated sites (n = more than 20, respectively) is shown in (F).

### Topographycal association of damaged DNA with RAD51 nuclear zones

Our findings suggest that dynamics of RAD51 is imposed to spatial constraints in the nucleus. Conversely, they imply that RAD51-mediated events during HR requires the transfer of DNAs into the bundle-like nuclear zone, but not the recruitment of RAD51 to DSB sites. To address this, we performed 3D-SIM analysis of the microirradiated nuclear areas. We examined the topographical association between phosphorylated RPA2 foci and bundle-like distributed EGFP-N-RAD51 signals. We noted a time-dependent increase of the correlation of EGFP-N-RAD51 with phosphorylated RPA2 foci after mIR (Fig 5A and C). This was confirmed by the detection of ssDNA with the immunostaining of bromodeoxyuridine (BrdU) incorporated into chromosomal DNA (Fig EV7). On the other hand, the correlation of EGFP-N-RAD51 with γH2AX signals did not increase (Fig 5B and C). This suggests that ssDNA formed outside the bundle-like nuclear zone, but not γH2AX labeled regions, shifts into such zones to enable repair by HR.

**Figure 5.**
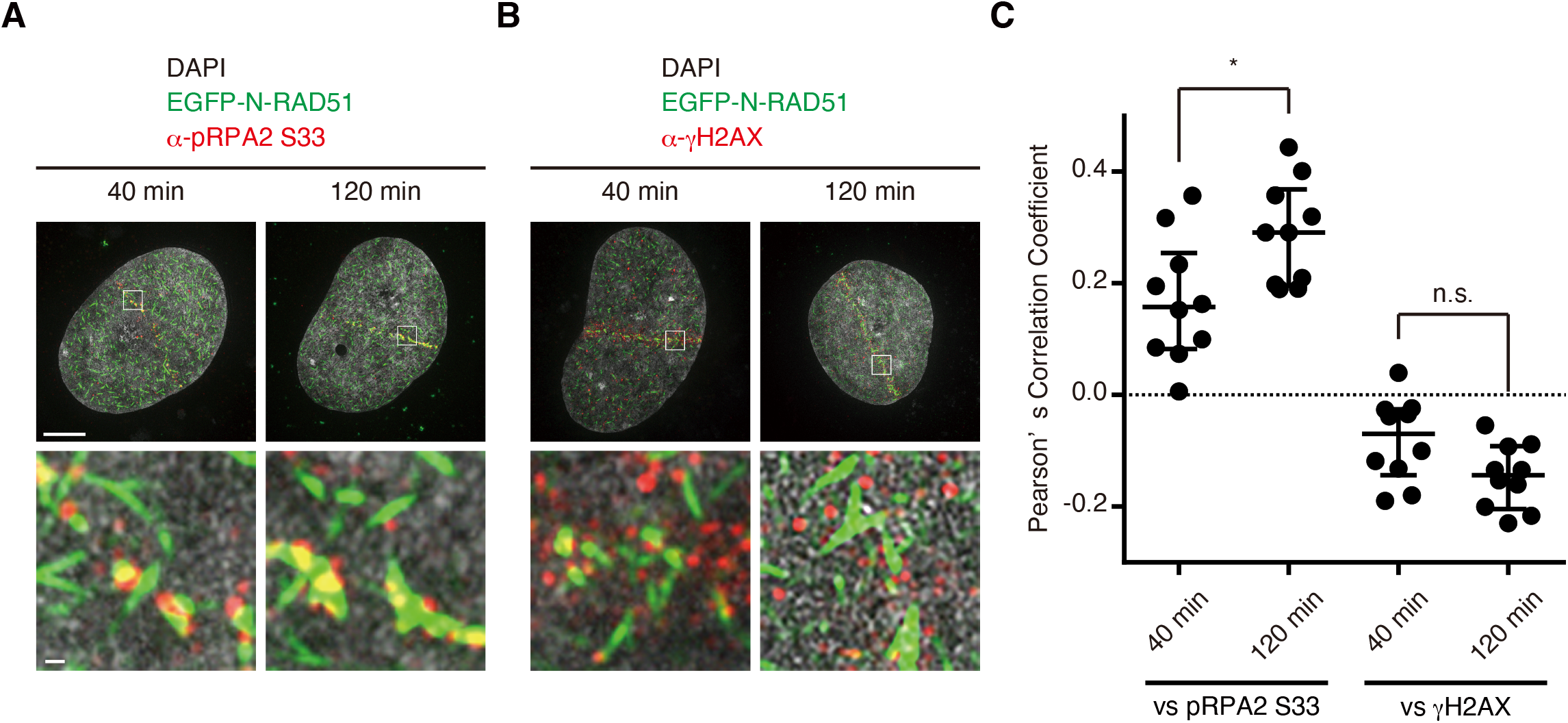
Elongated RAD51 foci provide distinctive DNA repair assemblies for HR. A,B Asynchronous GM0637 cells were transiently transfected with EGFP-N-RAD51 expression vector and incubated for 24 hours. Subsequently, UVA-microirradiation (mIR) was performed and fixed at 40 or 120 minutes after mIR. Phosphorylated RPA2 (S33) (pRPA2 S33) or γH2AX were detected by immunofluorescence and DNA was stained with DAPI. Z-stack images were acquired by 3D-SIM and representative projection images are shown. EGFP-N-RAD51, pRPA2 S33 (A) or γH2AX (B), and DNA were visualized in green, red and grayscale, respectively. Scale bars: 5 μm or 200 nm (insets). C Colocalization analysis based on voxel intensities with Pearson’s correlation coefficients. Data is presented as a Tukey dot-and-whiskers plot, with values that are above and below the whiskers drawn as individual dots. *P<0.05, ns = not significant, Brunner-Munzel test.

**Figure 6.**
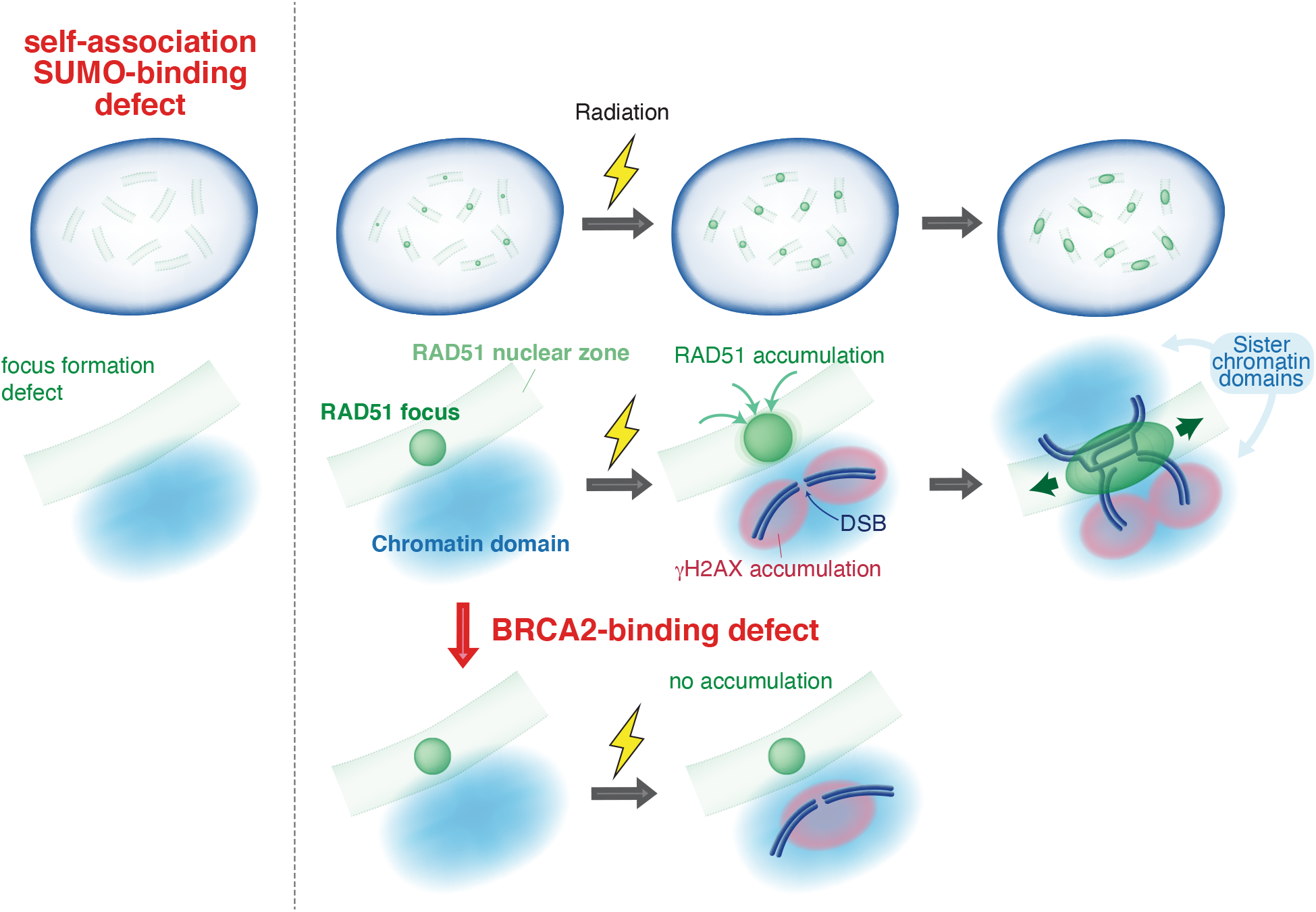
RAD51 repair assemblies. The scheme of RADA51 repair assemblies. RAD51 forms spherical foci for “storage” under normal conditions, dependent on self-association and the SUMO system. Following irradiation, more RAD51 accumulate adjacent to DSBs and elongate foci within the bundle-shaped distinctive nuclear zones for HR. This response to DNA damages requires the interaction of RAD51 with BRCA2. Thereafter, ssDNA generated for HR and its intact homologous template move into the “RAD51 repair assembly”, which provides a scaffold for homology search.

## Discussion

Current approaches have revealed the importance of the higher-order nuclear architecture, including dynamics of nuclear bodies, upon DNA metabolism. In the case of HR, when reflecting an open question of chromatin movements required for the pairing of damaged and intact sequences, one should take into account several, inherent problems of a higher order chromatin organization. With respect to the maintenance of the structural and functional integrity of 4D nucleome, the three-dimensional organization of nucleus in space and time, a minimal disruption of the 4D nucleome may provide a selective advantage.

In the present study, our analyses unveiled the existence of distinctive nuclear zones for HR. We found the elongation of RAD51 foci in a DSB-dependent manner (Fig 1). Notably, this “elongation”, not just swelling, implied the outline of such nuclear zones; this reflects the focus formation within these bundle-shaped areas. DAPI intensity classification assay further demonstrated that both endogenous RAD51 foci and the RAD51 bundle-like zones are localized within the identical classes, 3 to 5, independently of DNA damages (Fig 2 and 3). Moreover, there were no de novo RAD51 focus formation outside of the nuclear zone after microirradiation. Taken together, these findings suggest that RAD51 focus formation is strictly limited to distinctive nuclear zones.

That is, given that the range of motion of RAD51 is constrained only within the zone, transfer of damaged DNA outside of the zone is required for RAD51-mediated homologous pairing. In fact, our super-resolution microscopic analysis of RPA-coated ssDNA provided evidence supporting this idea (Fig 5A and C). This transfer was, intriguingly, not observed in the case of γH2AX (Fig 5B and C), suggesting a minimal disruption of chromatin structure. In order to exhibit an activity as a recombinase, the surface of RAD51 foci should face to decondensed chromatin loops, whereas illegitimate recombination reactions must be prevented. Considering this discrepancy, the distribution of RAD51 within relatively compact, not relaxed, region is quite a reasonable solution.

According to a recent study, a local increase in DSB movement is not a rate limiting for ectopic strand invasion during HR in yeast (Cheblal, Challa et al., 2020), suggesting that there may be an essential role played by the hierarchical levels of higher order chromatin folding in this recognition event. Therefore, macromolecular assemblies, such as RAD51 foci, may provide scaffold to facilitate homology search independent of local break movement. We argue that a particularly pronounced elongation of RAD51 foci in HR is required to bridge the spatial distance between sister chromatid domains (sCDs) which carry intact and broken DNA, respectively (Fig 2A, 3C and EV3B). This hypothesis is supported by evidence for the scaffolding properties of RecA protein, the *E. coli* homologue of RAD51 (Lesterlin, Ball et al., 2014, Wiktor, Gynnå et al., 2021). Importantly, RAD51 within the bundle-shaped nuclear zone exhibits increased mobility at damage sites in a BRCA2-dependent manner (Fig 4 and EV6B). Together with Fig 1, these observations suggest the presence of two types of RAD51 nuclear foci. One is foci for “storage”, and the other is those for “function”. The former shows spherical shape and static behavior; in contrast, the latter shows elongated shape and dynamic behavior. This static state of “storage” foci is quite reasonable to prevent unregulated recombination in S phase by sequestrating RAD51 in spite of the similar intranuclear distribution with “function” foci. Furthermore, it is conceivable that the difference in extent of elongation, indicated in Fig EV1, reflects the very spatial arrangement required for successful homology search. A strand containing a stalled replication fork is always located immediately next to its template strand within the same chromatin domain, allowing efficient RAD51-mediated strand invasion and fork restart (Zellweger, Dalcher et al., 2015). On the other hand, the distance between duplicated sister chromatid domains (sCDs) during G_2_ phase can be several hundred nanometers (Fig EV3F and G) (Stanyte et al., 2018).

The mechanistic details of how RAD51 foci elongate to form the unique bundle-shaped repair assemblies within the distinctive nuclear zones remains elusive. We expect that these assemblies harbor numerous other repair associated factors, even though the RAD51 recombinase is an essential and dominant component. In fact, the association of RAD51 with itself, with BRCA2 and with SUMO are all necessary for the elongated bundle formation, while its ATPase activity is not, intriguingly (Fig 4A and B). Since the ATPase activity is required for actual strand exchange, it follows that it is downstream from elongated bundle formation (Jackson & Durocher, 2013, West, 2003). Identifying the complex molecular details of specific structure-function relationships within the features of the nuclear landscape (Ochs, Karemore et al., 2019, Reindl, Girst et al., 2017, Strickfaden, Tolsma et al., 2020) will require a combination of top-down and bottom-up studies with integrated biophysical and biochemical approaches (Finn, Pegoraro et al., 2019, Marti-Renom, Almouzni et al., 2018, Szabo, Donjon et al., 2020, Takei, Yun et al., 2021).

## Materials and Methods

### Cell culture

The simian virus 40-transformed human fibroblast cell line GM0637 (Tashiro et al., 2000) and ataxia telangiectasia cell line AT5BIVA (and its ATM-proficient derivative 11-4) (Komatsu, Kodama et al., 1990, Nakada, 2005, Sun, Oma et al., 2010) were cultured in Dulbecco’s modified Eagle’s medium (DMEM) (Sigma-Aldrich), containing 10% fetal bovine serum (FBS) (JRH BioSciences). All cells were incubated at 37 °C in a humidified 5% (vol/vol) CO_2_ incubator.

### Plasmid Construction

The enhanced green fluorescent protein (EGFP) epitope-tagged RAD51 expression vectors were constructed as described previously (Shima et al., 2013). The transcription activator-like effector nuclease (TALEN) expression vectors were constructed as previously described (Ochiai, Miyamoto et al., 2014). Cells were transfected with the expression vectors except TALEN expression vectors, using either GeneJuice® (Millipore) or the FuGENE® HD (Promega) Transfection Reagent.

### Generation of DNA breaks

X-ray irradiation of cells was performed using the CP-160 cabinet X-radiator system (Faxitron). Hydroxyurea (Sigma-Aldrich) was used at a final concentration of 2 mM. TALEN expression vectors were transfected by the use of Lipofectamine® LTX reagent with PLUS reagent (ThermoFisher Scientific) and cells were incubated for 24 hours after transfection. UVA-microirradiation experiment (Walter, Cremer et al., 2003) was performed as described previously (Ikura, Tashiro et al., 2007); In brief, cells, in a 35 mm dish containing a 25 mm diameter coverslip (Matsunami), were transfected with 0.7-1.0 μg of EGFP-N-RAD51 expression vector. At 24 hours after transfection, Hoechst 33258 (Sigma-Aldrich) was added to culture medium at a final concentration of 2 μg/ml, to sensitize the cells for 10 minutes. For immunofluorescence staining using the anti-BrdU antibody, BrdU (Sigma-Aldrich) was added to culture medium at a final concentration of 10 μM and incubated in the dark for 24 hours before irradiation. Thereafter, culture medium was replaced by Leibovitz’s L-15 medium (Gibco), supplemented with 10% FBS, 25 mM HEPES (Gibco) and 200 nM Trolox (Sigma-Aldrich), and the cells were incubated at 37 °C for 10 minutes. The coverslips were subsequently transferred to a Chamlide TC live-cell chamber system (Live Cell Instrument) or Stage Top Incubator (INUBG2H-ELY, TOKAI HIT) mounted on the microscope stage, and maintained at 37 °C throughout the experiment. The UVA-microirradiation was performed using two confocal laser-scanning microscopes (Carl Zeiss Microscopy), and a Zeiss Laser Scan Microscope 780 (LSM780) confocal laser-scanning microscope with a 63x /1.4 C-apochromat objective was principally used. The 355 nm line of laser-UVA was used for microirradiation (three pulses at 570 μW). The 488 nm laser line was used for visualization of the EGFP fluorescence. Irradiated cells were incubated at 37 °C for the indicated periods of time. The other microscopy is a Zeiss Laser Scan Microscope 510 (LSM510) confocal laser scanning microscope with a 63x/1.4 plan-apochromat objective. Under this condition, the 364 nm line of the UVA-laser was used to introduce DBSs. The fluorescence recovery of EGFP-RAD51, bleached 3 times with a 488 nm HeNe-laser (0.5 mW) at 100% power, was monitored at 1% power, at the indicated time intervals.

### Immunofluorescence staining

Procedure for immunofluorescence staining has been previously described (Shima et al., 2013) and that for observation by three-dimensional structured illumination microscopy (3D-SIM) (Gustafsson, Shao et al., 2008) was adapted; Cells were rinsed with prewarmed (37 °C) PBS, and subsequently fixed with PBS containing 2% paraformaldehyde (PFA). In the case of pre-extraction, the cells were washed with prewarmed PBS and then incubated for 10 minutes on ice with CSK buffer (10 mM PIPES (pH 6.8), 100 mM NaCl, 300 mM sucrose and 3 mM MgCl_2_) containing 0.5% Triton X-100, 1 mM ethylene glycol bis (2-aminoethylether) tetraacetic acid (EGTA) and 1 mM phenylmethylsulfonyl fluoride (PMSF). Fixed cells were subsequently permeabilized with 0.5% Triton X-100 in PBS, and then washed three times with PBS. Before immunostaining, the coverslips were blocked with blocking solution. The coverslips were incubated at room temperature for 1 hour with the primary antibodies, and then washed three times with PBS. Thereafter, the coverslips were incubated with the appropriate goat secondary antibodies, conjugated to Alexa Fluor® 488, 568 or 594 fluorophores, at room temperature for 1 hour. After three washes with PBS, the nuclei were stained with PBS containing 2 μg/ml DAPI (Roche) for 10 minutes at room temperature. The coverslips were then washed with PBS three times, and the cells were subsequently postfixed using PBS containing 1.5% PFA, for 5 minutes at room temperature. After three washes with PBS, the samples were mounted in VECTASHIELD® (VECTOR LABORATORIES) antifade mounting medium and sealed with nail varnish.

### Fluorescence *in situ* hybridization on three-dimensionally preserved nuclei (3D-FISH) combined with immunostaining

FISH probes were purchased from Metasystems. 3D-FISH combined with immunostaining was performed as previously described (Solovei & Cremer, 2010). In brief, after transfection with TALEN expression vectors, AT5BIVA cells were fixed in PBS containing 4% PFA, and then permeabilized with 0.5% Triton X-100 in PBS. The coverslips were transferred into 20% glycerol/PBS and incubated for 60 min at room temperature. Next, these coverslips were frozen by liquid nitrogen and then thawed at room temperature. After six repeats of this freezing/thawing treatment, the samples were incubated in 0.1 N HCl. Then, these samples were equilibrated in 2x SSC and subsequently incubated in 50% formamide/2x SSC at 4°C overnight. The next day, 10 μl of probe mixture was loaded on a clean microscopic slide and the coverslips were placed on it. The samples were denatured, and then hybridized at 37°C overnight. After hybridization, the coverslips were washed in 0.4x SSC for 2min at 72°C and then incubated in 2x SSC containing 0.05% Tween-20 for 30 sec at room temperature. Immunostaining and postfixation follow brief rinse of coverslips in distilled water.

### Microscopy and image analysis

Image acquisition by conventional Widefield microscopy was performed using an Axioplan2 microscope equipped with an AxioCam MRm controlled by Axiovision (Carl Zeiss Microscopy). Confocal imaging was carried out on LSM 510 or LSM 780 microscopes. Live cell imaging and fluorescence recovery after photobleaching (FRAP) analyses were performed using a LSM510 or LSM780 confocal laser scanning microscope, as described previously (Ikura et al., 2007). The fluorescence recovery of EGFP-RAD51, bleached 3 times with a 488 nm HeNe-laser (LSM510) or Ar-laser (LSM780) at 100% power, was monitored at 1% (LSM510) or 0.1% (LSM780) power, at the indicated time intervals. The relative intensities in the bleached area were measured and normalized, using the average intensity before bleaching.

3D-SIM was mainly performed on a DeltaVision™ OMX microscope (GE Healthcare) version 4/Blaze equipped with PlanApo N 60x oil objective lens (numerical aperture of 1.42, Olympus). Different sCMOS cameras were used for each channel as the detector and the immersion oils with a refractive index of 1.512 or 1.514 were used.

DeltaVision™ OMX microscope version 3 equipped with a UPlanSApo 100x silicone oil objective lens (numerical aperture of 1.3, Olympus) and EMCCD cameras were also used. In this case, the same silicone oil was used. SI image stacks were acquired with a z-distance of 125 nm and with 15 raw SI images per plane (five phases and three angles). The SI raw datasets were computationally reconstructed with channel-specific measured optical transfer functions (OTFs) and a Wiener filter set between 0.001 and 0.006, using the softWoRx software package (Applied Precision). Adaptive image registration was performed by Chromagnon software (Demmerle, Innocent et al., 2017, Kraus, Miron et al., 2017, Matsuda, Koujin et al., 2020, Matsuda, Schermelleh et al., 2018); in short, we excited only the DAPI signal with a 405 nm laser and then acquired the signal in all required channels. The registration parameters were obtained from these DAPI images.

For scoring of intranuclear bundle-like distribution of EGFP-N-RAD51 mutants, images were obtained automatically with a CoolCube 1 camera, mounted with Metafer (MetaSystems). 200 EGFP positive cell nuclei per slide were scored.

### 3D image analysis using IMARIS® software

Following their acquisition, the 3D images were imported into IMARIS® 7.7 (Bitplane) for quantitative analyses by IMARIS® MeasurementPro 7.7 (Bitplane). The sphericity (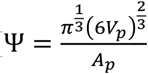; Vp = volume of the particle and Ap = surface area of the particle) (Wadell, 1935) was measured by isosurface rendering in IMARIS® 7.7 with a background subtraction algorithm and the following settings: experimentally defined threshold; noise filter eliminating structures <0.01 μm^3^ (Chapman, Sossick et al., 2012). Likewise, the Pearson’s correlation coefficients were calculated by IMARIS® Coloc 7.7 (Bitplane) to quantify the degree of colocalization between two fluorophores.

### Image analysis for DAPI density classification

Following their acquisition, the images were converted to 16-bit composite TIFF stacks, using softWoRx® Explorer (Applied Precision). Thereafter, the DAPI intensity quantification was performed as previously described (Schmid et al., 2017); in brief, DAPI-stained DNA was divided into seven classes with equal intensity variances. The open-source statistics software R (R Project for Statistical Computing) was used for the analysis.

### Quantitative PCR assay

For detection of the cleavage efficiency of TALEN, quantitative PCR was performed with 5 ng purified genomic DNA as the template, using SYBR® *Premix Ex Taq*™ (Tli RNaseH Plus) (Takara Bio), LightCycler® FastStart DNA Master SYBR GREEN I (Roche) and a LightCycler® II real-time PCR system (Roche). Dissociation curve analysis of the amplified DNA melting temperature showed that each primer set gave a single and specific product (Sun et al., 2010). Primers for real-time PCR were

bt56 forward: 5’TACTCTGAATCTCCCGCA3’
bt56 reverse: 5’CGCTCGTTCTCCTCTAA3’

### *In vitro* D-loop assay

Human RAD51 and human RAD51-GFP were prepared by the methods described previously (Ishida, Takizawa et al., 2008, Kobayashi, Sekine et al., 2014). For the GFP-RAD51 protein, the stop codon of the enhanced *GFP* (*EGFP*) gene was replaced by a DNA fragment encoding Ser-Gly-Leu-Arg-Ser residues, as a linker peptide, and human RAD51 gene was fused after the linker peptide sequence. The resulting *GFP-RAD51* gene was ligated into the *Nde*I-*Bam*HI site of the pET15b vector (Novagen). Human GFP-RAD51 was purified by the same method as that for human RAD51-GFP, except a washing step with the buffer containing 100 mM KCl for Heparin Sepharose column chromatography (GE Healthcare). For the D-loop formation assay, high-pressure liquid chromatography-purified deoxyribo-oligonucleotide was purchased from Nihon Gene Research Laboratory: 5S 70-mer ssDNA, 5’-CCGGT ATATT CAGCA TGGTA TGGTC GTAGG CTCTT GCTTG ATGAA AGTTA AGCTA TTTAA AGGGT CAGGG-3’ The superhelical double-stranded DNA containing tandem 5S rDNA repeats was prepared as described previously (Kagawa, Kurumizaka et al., 2001). The D-loop formation assay was performed as described previously (Kobayashi et al., 2014). The reactions were conducted in reaction buffer containing 24 mM HEPES (pH 7.5), 30 mM NaCl, 1 mM ATP, 20 mM creatine phosphate, 75 μg/ml creatine kinase, 1 mM DTT, 1 mM MgCl_2_, 2 mM CaCl_2_, 4% glycerol, and 100 μg/ml BSA. The indicated amount of RAD51, GFP-RAD51, or RAD51-GFP was incubated with the ^32^P-labeled 5S ssDNA 70-mer (1 μM in nucleotides) at 37°C for 10 min. The reactions were then initiated by the addition of the supercoiled dsDNA (30 μM in nucleotides), and were continued at 37°C for indicated times. The homologous pairing products, D-loops, were detected as described previously (Kobayashi et al., 2014).

### Western blotting analysis

Whole cell extracts were resolved by sodium dodecyl sulfate-polyacrylamide gel electrophoresis (SDS-PAGE) on SuperSep™ Ace 5-20% gradient gels (Wako Pure Chemical Industries) and electrotransferred to nitrocellulose membranes (Pall Corporation). Membranes were blocked in Blocking One (Nacalai Tesque) solution and subsequently incubated with primary antibodies diluted in Blocking One solution for 1 hour at room temperature. After four washes in Tris-buffered saline with Tween20 (TBS-T), the membranes were incubated with horseradish peroxidase-conjugated secondary antibodies, diluted in Blocking One solution for 1 hour at room temperature. The membranes were then washed in TBS-T four times, and treated with Luminata™ Forte Western HRP Substrate (Millipore).

### Statistics and reproducibility

The non-parametric Mann–Whitney-Wilcoxon test was performed with KaleidaGraph Version 4.1 (Synergy Software). The non-parametric Brunner-Munzel tests were performed with R. The generation of plots were performed with GraphPad Prism 6 or 8 (GraphPad Software). Experiments were not randomized and no blinding was used during data analysis. Sample size, statistical tests and the number of biological replicates for each experiment are indicated in the figure legends.

### Antibodies

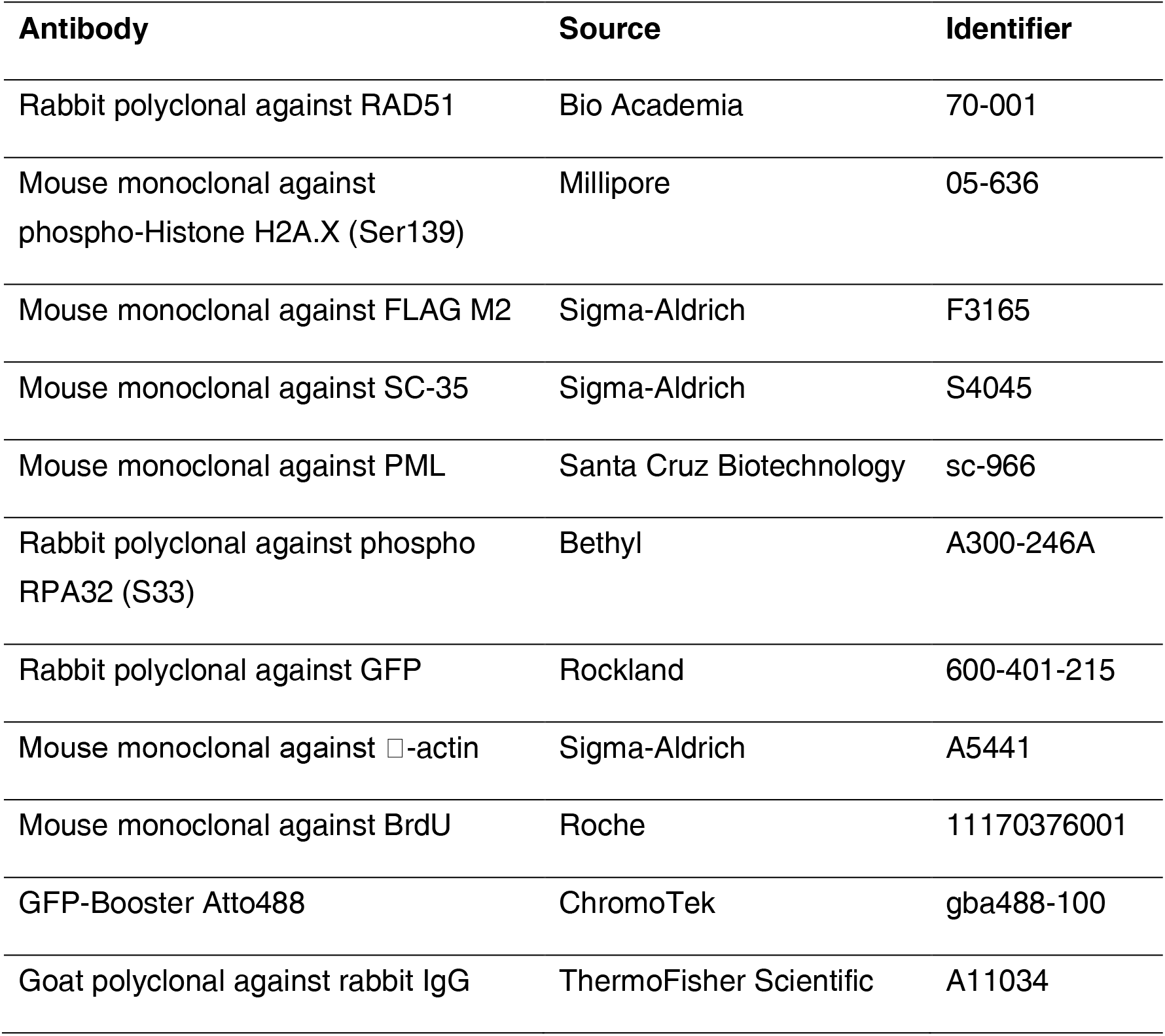
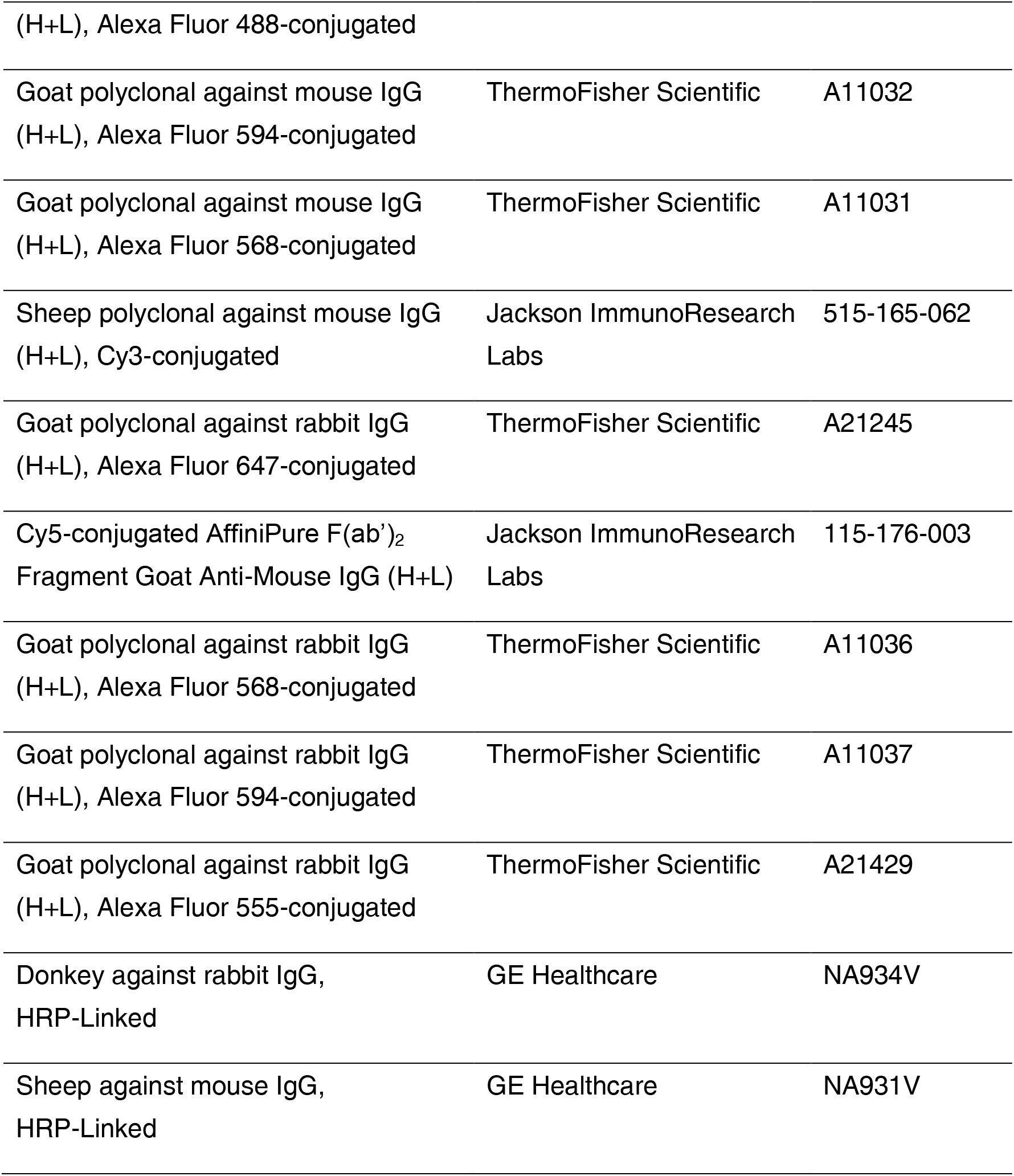

## Acknowledgements

We thank Drs. Atsushi Matsuda, Lothar Schermelleh and Yasuhiro Hirano for technical assistance with the 3D-SIM, and Ms. Aiko Kinomura for technical assistance with 3D-FISH. We also thank Heinrich Leonhardt and members of the Center of Advance Light Microscopy (CALM) of the LMU Biocenter for their support in initial 3D-SIM studies. We thank Drs. Yasushi Hiraoka, Tokuko Haraguchi, Christoph Cremer, Marion Cremer, Takehiko Shibata, Shun Matsuda, Tomonari Matsuda, Friedl Anna, all current and former members of the S. Tashiro laboratory and RcMcD, especially Dr. Kazuteru Kono and Mr. Hidekazu Suzuki, for helpful discussions. This study was supported by JSPS KAKENHI Grants (Numbers JP18K18195 to Y. H., JP26430114 and JP19K07641 to J. S., JP21K15017 to Y. K., JP15H02506 to K. I. and JP15H02821, JP16H01312, JP18H03373 and JP21H04927 to S. Tashiro) and the Platform Project for Supporting in Drug Discovery and Life Science Research (Platform for Dynamic Approaches to Living System) from the Ministry of Education, Culture, Sports, Science and Technology (MEXT) and Japan Agency for Medical Research and Development (AMED). This work was partially supported by JST CREST (JPMJCR18S3), the International Atomic Energy Agency (IAEA CRP E3.50.08), the Program of the network-type joint Usage/Research Center for Radiation Disaster Medical Science of Hiroshima University, Nagasaki University and Fukushima Medical University, and Core Research for Organelle Diseases in Hiroshima University (the MEXT program for promoting the enhancement of research universities, Japan).

## Author contributions

T.C. and S.Tashiro proposed the project. H.S. led the test experiments. Y.H. and S.Tashiro designed the experimental setup. W.K. performed the in vitro D-loop formation assay with supervision by H.K.. V.J.S. performed the quantitative analyses of DAPI intensity classes with Y.H.. H.O. constructed TALEN expression vectors, and J.S. determined its optimal experimental condition with Y.H.. Y.M. performed initial 3D-SIM microscopy of EGFP-N-RAD51 expressing cells. Y.H. performed all other experiments, collected data, analyzed and interpreted results with S. Tashiro. H.K., T.I., S. Tate, and K.I. supervised research. Y.H., T.C., and S. Tashiro drafted the manuscript. All authors read the manuscript, gave comments, suggested changes, and agreed with the final version.

## Expanded View Figure legends

**Figure EV1. Function-dependent dynamic shaping of RAD51 foci**

Before and after exposure to 2 Gy X-ray or 2 mM hydroxyurea (HU), asynchronous GM0637 cells were pre-treated with CSK buffer and then fixed. X-ray-exposed cells were fixed at 2 hours after irradiation and HU-exposed cells were fixed at indicated time points after treatment. Endogenous RAD51 and γH2AX were detected by immunofluorescence and DNA was stained with DAPI. Z-stack images were acquired by 3D-SIM. Statistical analysis of the sphericities of the respective RAD51 foci in cell nuclei under indicated experimental conditions. Data is presented as a Tukey box-and-whiskers plot, with values that are above and below the whiskers drawn as individual dots. ***P<0.0001, Brunner-Munzel test.

**Figure EV2. DSB induction by use of TALEN system**

A Schematic representation of the hybridization probe targeting region for fluorescence in situ hybridization on three-dimensionally preserved nuclei (3D-FISH) and the site designed for cleavage by TALEN-L and TALEN-R.

B Asynchronous AT5BIVA cells were transiently transfected with TALEN-L and TALEN-R expression vectors and incubated for indicated time points. Efficiency of cleavage of the TALEN-targeted sequence was evaluated by quantitative PCR assay, using primers shown in (A).

C Asynchronous AT5BIVA cells were transiently transfected with TALEN-L and TALEN-R expression vectors and fixed at 24 hours after transfection. FLAG-tagged TALEN proteins were detected by indirect immunofluorescence staining using anti-FLAG antibody and DNA was stained with DAPI. A Representative image acquired by wide-field microscopy is shown. In merged image, TALEN proteins and DNA were visualized in red and blue, respectively. Scale bar: 5 μm.

**Figure EV3. 3D-FISH in combination with TALEN system**

A Outline of the experimental procedure.

B Representative confocal images of MLL locus with RAD51 focus in AT5BIVA cell nuclei, detected by fluorescence in situ hybridization on three-dimensionally preserved nuclei (3D-FISH) in combination with immunofluorescence staining using anti-RAD51 antibody. DNA was stained with DAPI. In the top panels, the respective signals were indicated in grayscale. In the middle panels, each FISH probe was indicated in green or red and RAD51 signals were indicated in grayscale. In the bottom panels, FISH probes were indicated in grayscale and the boundaries of RAD51 signals were indicated by dashed line. Scale bar: 200 nm.

C Percentage of MLL locus with RAD51 signal in TALEN-treated cell nuclei. L: cells expressing only TALEN-L, R; cells expressing only TALEN-R, L+R: cells expressing both TALEN-L and -R.

D Representative confocal image of MLL loci with RAD51 foci. Each MLL locus was further classified into two groups according to the distance between respective gravity centers of red and green signal intensities: adjacent (< 1 μm) or separated (> 1 μm apart). Scale bar: 1 μm.

E The Venn diagrams depict MLL loci with adjacent red/green signals and with RAD51 signal (n = 38 sister chromatids/19 independent cells).

F, G 3D spatial distance between duplicated MLL gene loci on 11q23. Representative images acquired by 3D-SIM are shown in (F). DNA was stained with DAPI and visualized in grayscale. Scale bar: 200 nm. 3D distance between green and red signal pairs was measured in nuclei with “two distinct green or red signals”. Data is presented as a Tukey dot-and-whiskers plot, with values that are above and below the whiskers drawn as individual dots.

**Figure EV4. Distribution of non-chromatin nuclear domains**

A Asynchronous GM0637 cells were transiently transfected with EGFP-N-RAD51 expression vector and fixed at 24 hours after transfection. Non-chromatin nuclear domains, nuclear speckle (SC35) or PML body, were detected by immunofluorescence and DNA was stained with DAPI. Z-stack images were acquired by 3D-SIM and a representative projection image is shown. EGFP-N-RAD51, SC35 and DNA were visualized in green, red and grayscale, respectively. Scale bars: 5 μm and 200 nm (insets).

B The over/underrepresentation of nuclear speckle (black) or PML body (gray) in DAPI intensity classes of GM0637 cell nuclei (n = 21), ranging from class 1 (close to background intensity) to class 7 (highest). Values represent means with SD.

**Figure EV5. Functional assays for EGFP-fused RAD51**

A Purified human RAD51 (0.75 μg), GFP-RAD51 (0.75 μg), and RAD51-GFP (0.75 μg) were analyzed by 12% SDS-PAGE with Coomassie Brilliant Blue staining. Lane 1 indicates the molecular mass markers.

B The scheme of D-loop formation assay.

C, D RAD51 (400 nM), GFP-RAD51 (400 nM), or RAD51-GFP (400 nM) was incubated with ^32^P-labeled 5S 70-mer ssDNA (1 μM in nucleotides). The reactions were initiated by addition of superhelical dsDNA (30 μM in nucleotides), and were continued at 37°C for 5, 10, and 20 min. The average values of three independent experiments are shown in (D), with the standard deviation values.

E Asynchronous GM0637 cells were transiently transfected with EGFP-N-RAD51 expression vector and incubated for 24 hours. Subsequently, fluorescence recovery after photobleaching (FRAP) analysis of EGFP-N-RAD51 in combination with UVA-microirradiation (mIR) was performed; the area enclosed by dashed line was bleached. Images were acquired at the indicated time points. Scale bar: 5 μm.

**Figure EV6. Expression levels of EGFP-N-RAD51 mutants and dynamics of EGFP-N-RAD51-WT at UVA-irradiated or unirradiated areas**

A Asynchronous GM0637 cells were transiently transfected with either one of the indicated types of mutant EGFP-N-RAD51 expression vectors, defective in self-association (F86E), ATPase activity (G151D), BRCA2 binding (A190/192L) or interaction with SUMO (V264K), respectively. Cells were harvested at 24 hours after transfection and an immunoblotting analysis of the respective cell extracts using anti-GFP and anti-RAD51 antibodies was performed. The immunoblots was also developed using an anti-β-actin antibody as a loading control.

B Asynchronous GM0637 cells were transiently transfected with EGFP-N-RAD51 expression vector and incubated for 24 hours. Subsequently, fluorescence recovery after photobleaching (FRAP) analysis of EGFP-N-RAD51 in combination with UVA-microirradiation (mIR) was performed: stripe-type microirradiation (pink arrow in “MI”). Images acquired at the indicated time points were indicated in the left panel: the area enclosed by dashed line was bleached. Scale bar: 5 μm. Profile plots of the respective signal intensities along the arrows, measured at each time point indicated, are shown in the right panel.

**Figure EV7. Topographical association between single stranded DNA and bundle-like distributed EGFP-N-RAD51 signals**

A Asynchronous GM0637 cells were transiently transfected with EGFP-N-RAD51 expression vector and incubated in culture medium containing 10 μM bromodeoxyuridine (BrdU) for 24 hours. Subsequently, UVA-microirradiation (mIR) was performed and fixed at 120 minutes after mIR. BrdU was detected by immunofluorescence and DNA was stained with Hoechst 33342: An anti-BrdU antibody was used to detect single stranded DNA induced by UVA mIR. Representative images acquired by wide-field microscopy are shown. In merged images, EGFP-N-RAD51 and BrdU and DNA were visualized in green and red, respectively. Scale bar: 5 μm.

B Asynchronous GM0637 cells were transiently transfected with EGFP-N-RAD51 expression vector and incubated in culture medium containing 10 μM BrdU for 24 hours. Subsequently, UVA-mIR was performed and fixed at 120 minutes after mIR. BrdU was detected by immunofluorescence and DNA was stained with DAPI. Z-stack images were acquired by 3D-SIM. Representative projection images were shown. In merged images, EGFP-N-RAD51 and BrdU were visualized in green and red, respectively. Scale bars: 5 μm or 200 nm (insets).

C Asynchronous GM0637 cells were transiently transfected with EGFP-N-RAD51 expression vector and incubated in culture medium containing 10 μM BrdU for 24 hours. Subsequently, UVA-mIR was performed and fixed at 30 or 90 minutes after mIR. BrdU and phosphorylated RPA2 (S33) (p-RPA2 S33) were detected by immunofluorescence and DNA was stained with Hoechst 33342. Representative images acquired by wide-field microscopy are shown. In merged images, EGFP-N-RAD51, BrdU and p-RPA2 were visualized in green, blue and red, respectively. Scale bar: 5 μm.

